# Structural basis for human T-cell leukemia virus type 1 Gag targeting to the plasma membrane for assembly

**DOI:** 10.1101/2021.04.08.439007

**Authors:** Dominik Herrmann, Lynne W. Zhou, Heather M. Hanson, Nora A. Willkomm, Louis M. Mansky, Jamil S. Saad

**Affiliations:** Department of Microbiology, University of Alabama at Birmingham, Birmingham, AL 35294; Institute for Molecular Virology, University of Minnesota – Twin Cities, Minneapolis, MN 55455 USA

**Keywords:** Human T-cell leukemia virus type 1 (HTLV-1), human immunodeficiency virus type 1 (HIV-1), retrovirus, virus assembly, Gag polyprotein, myristoylated matrix (MA), plasma membrane (PM), phosphatidylinositol 4,5-bisphosphate (PI(4,5)P_2_), inositol hexakisphosphate (IP_6_), inositol trisphosphate (IP_3_), phosphatidylcholine (PC), phosphatidylserine (PS), liposome, nuclear magnetic resonance (NMR), isothermal titration calorimetry (ITC)

## Abstract

During the late phase of retroviral replication, the virally encoded Gag polyprotein is targeted to the plasma membrane (PM) for assembly. Gag–PM binding is mediated by the N-terminal matrix (MA) domain of Gag. For many retroviruses, Gag binding to the PM was found to be dependent on phosphatidylinositol 4,5-bisphosphate [PI(4,5)P_2_]. However, it was shown that for human T-cell leukemia virus type 1 (HTLV-1), Gag binding to membranes is less dependent on PI(4,5)P_2_, suggesting that other factors may modulate Gag assembly. To elucidate the mechanism by which HTLV-1 Gag binds to the PM, we employed NMR techniques to solve the structure of unmyristoylated MA (myr(–)MA) and to characterize its interactions with lipids and liposomes. The MA structure consists of four α-helices and unstructured N- and C-termini. We show that myr(–)MA binds to PI(4,5)P_2_ via the polar head and that myr(–)MA binding to inositol phosphates (IPs) is significantly enhanced by increasing the number of phosphate groups on the inositol ring, indicating that the MA–IP binding is governed by charge–charge interactions. The IP binding site was mapped to a well-defined basic patch formed by lysine and arginine residues. Using a sensitive NMR-based liposome binding assay, we show that myr(–)MA binding to membranes is significantly enhanced by phosphatidylserine (PS). Confocal microscopy data show that Gag is localized to the inner leaflet of the PM of infected cells, while the Gag G2A mutant, lacking myristoylation, is diffuse and cytoplasmic. These findings advance our understanding of a key mechanism in retroviral assembly.

---

During the late phase of retroviral replication, the virally encoded Gag polyproteins are targeted to the plasma membrane (PM) for assembly, virus budding and release (1-12). During or subsequent to virus budding, the virally encoded protease cleaves off Gag protein into matrix (MA), capsid (CA), nucleocapsid (NC), and short peptides to form mature virions (reviewed in (9,13,14)). It is demonstrated that Gag binding to the PM is mediated by the MA domain, which for most retroviruses contains a bipartite signal consisting of an N-terminal myristoyl (myr) group and a highly basic region. Over the last three decades, studies have established that binding of retroviral Gag to membranes is regulated by many factors such as protein multimerization, cellular and viral RNA, and the type of lipids and degree of acyl chain saturation (6,7,15-33).

HIV-1 Gag binding to the PM was shown to be dependent on phosphatidylinositol 4,5-bisphosphate [PI(4,5)P_2_] (7), a PM component that fulfills many cellular functions by acting as a substrate for numerous proteins (34,35). Over-expression of phosphoinositide 5-phosphatase IV (5ptaseIV), which reduces PI(4,5)P_2_ levels by hydrolyzing the phosphate at the D5 position of PI(4,5)P_2_, led to significant reduction in Gag–PM localization and attenuation of virus production (7). PI(4,5)P_2_–dependent Gag assembly has also been shown for other retroviruses such as HIV-2 (12), Mason-Pfizer monkey virus (MPMV) (36,37), murine leukemia virus (10), feline immunodeficiency virus (38), and avian sarcoma virus (39-41).

NMR structural studies of HIV-1 MA binding to PI(4,5)P_2_ containing truncated (*tr*) acyl chains have shown *tr*-PI(4,5)P_2_ binding to MA induced a conformational change that promoted myr exposure (42). The structure of MA–*tr*-PI(4,5)P_2_ complex showed that both the polar head and the truncated 2’-acyl chain are involved in binding (42). Subsequent studies have shown that PM lipids such as PS, PC, and phosphatidylethanolamine (PE) with truncated acyl chains also bind to HIV-1 MA (43). NMR studies confirmed MA binding to membrane mimetics such as bicelles, micelles and lipid nanodiscs (43,44). NMR studies of HIV-2 (12) and MPMV MA proteins (37) as well as surface plasmon resonance studies of HIV-1 Gag and/or MA proteins (45) have shown that MA proteins are capable of interacting with the acyl chains of phosphoinositides, and that increasing the length of the acyl chain resulted in stronger binding. Employing computational methods (46) and an NMR-based liposome assay (47), it was suggested that acyl chains of native PI(4,5)P_2_ are not involved in MA binding and that Gag– membrane interaction is mediated predominantly by dynamic, electrostatic interactions between conserved basic residues of MA and PI(4,5)P_2_/PS (46,47).

Most recent cryo-electron tomography data revealed that MA undergoes dramatic structural maturation to form very different lattices in immature and mature HIV-1 particles (48). Mature MA forms a hexameric lattice in which the acyl chain of a phospholipid extends out of the membrane to bind a pocket in MA, consistent with the NMR studies (42). Based on these studies, it was suggested that maturation of HIV-

1 not only achieves assembly of the capsid surrounding the RNA genome, but it also extends to repurpose the MA lattice for an entry or post-entry function and causes partial removal of 2,500 acyl chains from the viral membrane (48). Taken together, despite some differences in the proposed models, these studies have shed new insights on how various retroviral Gag proteins interact with the inner leaflet of the PM.

In this report, we focus on the molecular mechanism by which human T-cell leukemia virus type 1 (HTLV-1) Gag polyproteins are targeted to the PM for assembly. HTLV is a zoonotic virus with simian T-cell leukemia virus counterparts found in monkeys. HTLV-1 and HTLV-2 are the most studied subtypes of HTLV. Even though they share ∼ 70% nucleotide identity and have a similar genome structure, HTLV-1 is considered more pathogenic as it is associated with adult T-cell leukemia and HTLV-1 associated myelopathy/tropical spastic paraparesis (49-51). HTLV-1 transmission occurs mainly through cell-to-cell contacts rather than cell-free virus particles (52,53). In addition, HTLV-1 infected T-cells can multiply by clonal expansion, consequently increasing the viral burden without the need for virus replication and reinfection (54,55). In order to develop effective antiretroviral treatments for HTLV-1, it is paramount to gain a more complete understanding of the molecular processes that govern HTLV-1 replication. However, many fundamental aspects of HTLV-1 replication, including particle assembly, are incompletely understood.

It has been shown that HTLV-1 Gag binding to the PM and to liposomes is less dependent on PI(4,5)P_2_ (30). Unlike HIV-1 Gag, subcellular localization of and VLP release by HTLV-1 Gag were minimally sensitive to 5ptaseIV overexpression, suggesting that the interaction of HTLV-1 MA with PI(4,5)P_2_ is not a key determinant for HTLV-1 particle assembly (30,31). It was also shown that although PI(4,5)P_2_ enhanced HTLV-1 Gag binding to liposomes, Gag proteins bound efficiently to liposomes lacking PI(4,5)P_2_ but containing phosphatidylserine (PS) if similar overall negative charge is maintained (30). HTLV-1 Gag was found to bind to membranes with a higher affinity than that of HIV-1 Gag with no susceptibility to RNA-mediated inhibition of membrane binding (30). Chimeric switching of MA domains between HIV-1 and HTLV-1 Gag proteins showed that these differences are mediated by the MA domain of Gag (30). Subsequent studies have shown that single amino acid substitutions that confer a large basic patch rendered HTLV-1 MA susceptible to the RNA-mediated block, suggesting that RNA blocks MA containing a large basic patch (31). These data supported a model in which HTLV-1 Gag localizes to the PM via the MA domain with higher efficiency but less specificity than for other retroviruses (30,31).

Further comparison of the subcellular localization of HIV-1 with HTLV-1 Gag *in vivo* using dual-color, z-scan fluorescence fluctuation spectroscopy and total internal reflection fluorescence microscopy revealed significant differences in the cytoplasmic threshold concentration of Gag required for PM binding (56). Cytoplasmic HTLV-1 Gag associated with the PM at nanomolar concentrations, whereas HIV-1 Gag approached μM concentrations before it was observed at the PM. This dramatic difference in binding affinity highlights the need for a sharper understanding of retroviral Gag localization to the PM on the molecular level.

Herein, we characterized the interactions of HTLV-1 unmyristoylated MA protein (myr(–)MA) with lipids and liposomes by NMR and isothermal titration calorimetry (ITC) methods. Our data revealed that MA contains a PI(4,5)P_2_ binding site and that myr(–)MA binding to membranes is enhanced significantly by phosphatidylserine (PS). Confocal microscopy data show that Gag is localized to the inner leaflet of the PM, while the Gag G2A mutant lacking myristoylation is diffuse and cytoplasmic, leading to severe attenuation of particle production. These findings advance our understanding of a key mechanism in retroviral assembly.

## Results

### Structure determination of HTLV-1 myr(–)MA

The HTLV-1 MA domain consists of 130 residues and is naturally myristoylated. Due to technical challenges, we were unable to produce soluble, homogenous and monodisperse myristoylated HTLV-1 MA via recombinant techniques. Therefore, our studies were all conducted with HTLV-1 myr(–)MA. First, we generated a structural model of HTLV-1 myr(–)MA using I-TASSER (57,58). As expected, the resulting structural model indicated that the proline-rich C-terminal domain lacks an ordered structure. This model is consistent with the structural data of the closely related HTLV-2 myr(–)MA protein (59). Therefore, we truncated the C-terminal tail of MA (residue 100-130) to generate MA_99_ for expression via recombinant techniques. Solution properties of full-length myr(–)MA and myr(–)MA_99_ proteins were analyzed by a gel filtration mobility assay (Fig. S1). A gel filtration mobility assay with known protein standards revealed that the estimated molecular weight of myr(–)MA and myr(–)MA_99_ proteins are ∼24 and 10 kDa, respectively (Fig. S1). Whereas the estimated molecular weight of myr(–)MA appears to be higher than the calculated monomeric unit (∼15 kDa), no evidence for protein self-association was observed at all tested protein concentrations. The migration behavior of myr(–)MA is likely attributed to its shape caused by the unstructured C-terminal 31 residues. A minor species (∼10 %) of disulfide cross-linked dimer via Cys^61^ was observed during purification and was eliminated by inclusion of TCEP in buffers.

2D ^1^H-^15^N HSQC data obtained for myr(–)MA and myr(–)MA_99_ confirmed that truncation of the C-terminal 31 residues did not adversely affect the structure and/or fold of the globular domain (Figs. 1A and S2). Standard triple-resonance, NOESY, and TOCSY experiments were collected for HTLV-1 myr(–)MA_99_, which were used to generate near-complete backbone and side-chain chemical shift assignments. Subsequently, an initial list of distance restraints was created using Unio’10 Atnos/Candid functionality of automated, iterative peak picking of raw NOESY spectra. The list of distance constrains was extended by manual analysis of NOE-based spectra, and structures were calculated using CYANA. Superposition of the 20 lowest-penalty myr(–)MA_99_ structures is shown in Figure S3 (see also Table S1). The globular domain of myr(–)MA_99_ extends from residue 21 to 93 and consists of four α-helices, similar to that observed for HTLV-2 myr(–)MA (Figs. 1B and S4) (59). Not surprisingly, because HTLV-1 and HTLV-2 MA proteins share ∼60% sequence identity, their structures exhibit an overall similar fold. However, some differences were observed in helix packing and orientations (Fig. S4). Notably, we identified numerous unambiguous NOEs between residues Phe^27^ and Leu^84^/Leu^87^, and Tyr^65^ and Ile^63^/Leu^87^/Gln^91, 4 2^ resulting in a tight packing of helix III against helix IV (Fig. 2). Additionally, NMR data did not support the existence of a stable 3_10_ helix, previously observed between helices II and III in HTLV-2 myr(–)MA (59). The myr(–)MA_99_ structure exhibits a well-defined right-handed turn in this area but lacks the critical amide (*i, i+3*) or amide (*i, i+4*) NOEs, indicating that this region is more flexible than in HTLV-2 myr(–)MA. Rapid solvent-exchange of the amide proton of His^22^ precluded its assignment, as it was the case in HTLV-2 myr(–)MA (59). However, using cross-peak patterns in other spectra we were able to assign the carbon and proton chemical shifts for His^22^. Determination of the three-dimensional structure of HTLV-1 MA_99_ is a necessary step for characterizing MA–lipid and MA–membrane interactions.

**Figure 1.**
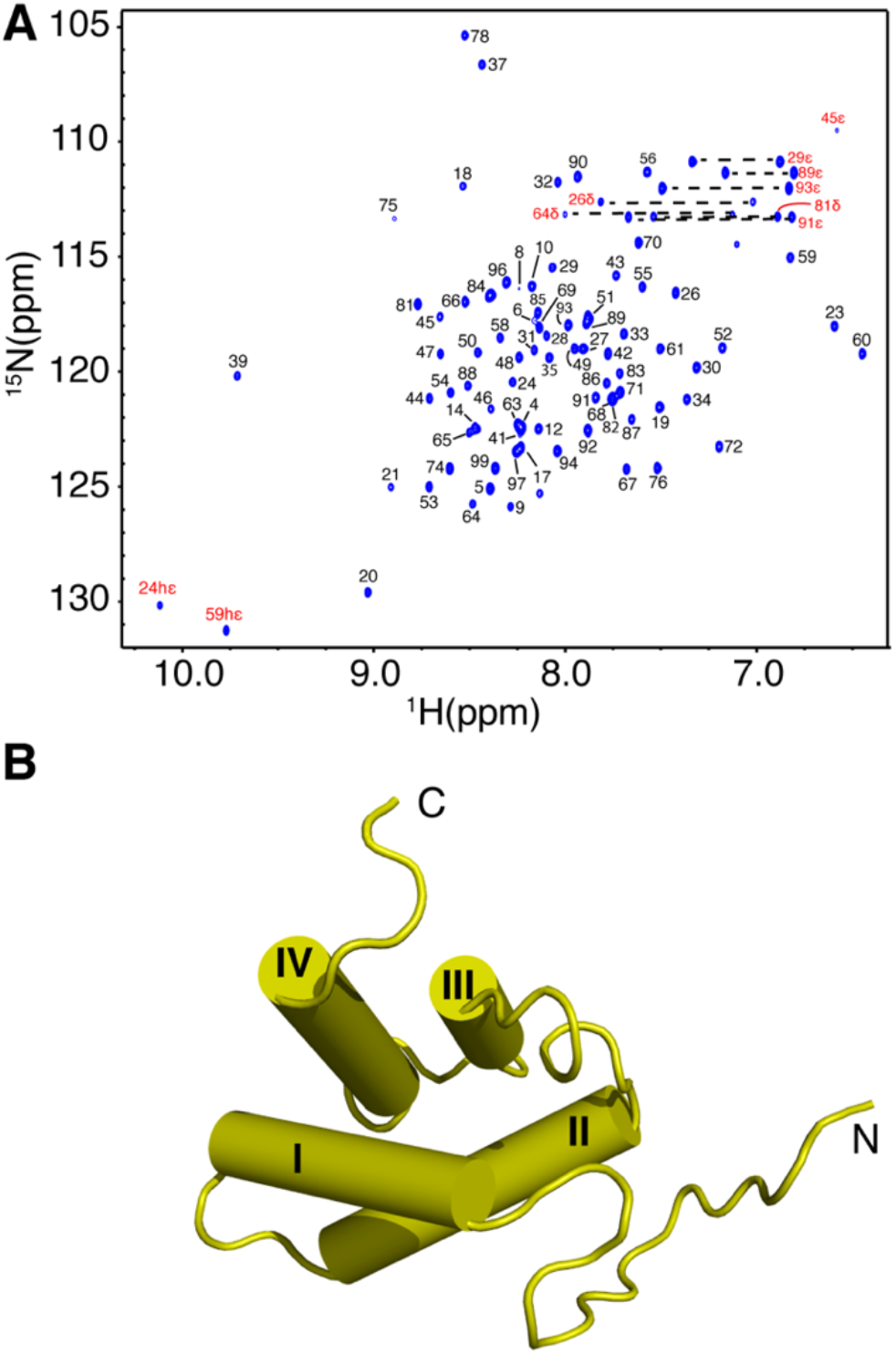
NMR data and structure of HTLV-1 myr(–)MA_99_. *A*, 2D ^1^H-^15^N HSQC NMR spectrum of the myr(–)MA_99_ protein at 35 °C in 50 mM sodium phosphates (pH 6.5), 100 mM NaCl and 2 mM TCEP. *B*, Cartoon representation of the myr(–)MA_99_ structure.

**Figure 2.**
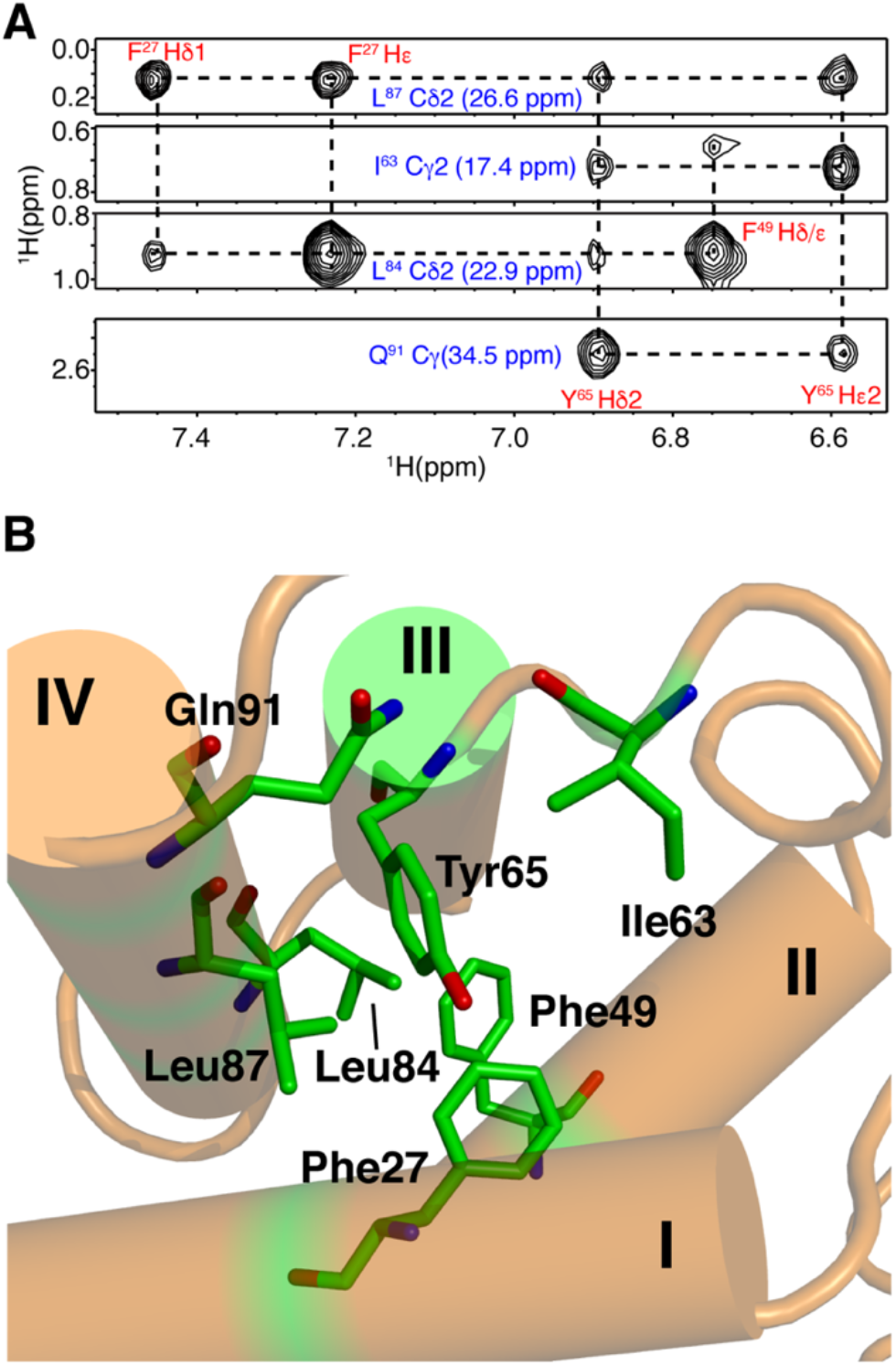
NMR data and structure of HTLV-1 myr(–)MA_99_. *A*, Selected ^1^H–^1^H strips from the ^13^C-edited HMQC-NOESY spectrum showing unambiguous NOEs between residues on helices III and IV. *B*, Cartoon and stick structural view showing relationship between residues represented by the NOEs above.

### PI(4,5)P_2_ binding to HTLV-1 myr(–)MA

Native PI(4,5)P_2_ has a high propensity to form micelles in aqueous solution (60), which causes severe signal broadening in the NMR spectra as described in our earlier studies (12,42). Therefore, we used dibutanoyl-PI(4,5)P_2_ [diC_4_-PI(4,5)P_2_], a soluble analog with truncated acyl chains (see Fig. S5 for chemical structures of lipids used in this study). This ligand was extensively used in previous studies of retroviral MA proteins (11,12,37,40,42,45,61). Due to the somewhat higher propensity of myr(–)MA_99_ protein to precipitate out at high lipid concentrations, which precluded accurate measurement of binding parameters, all studies below were conducted with the full-length myr(–)MA protein.

Titration of HTLV-1 myr(–)MA with increasing amounts of diC_4_-PI(4,5)P_2_ led to significant chemical shift perturbations (CSPs) for a subset of ^1^H and ^15^N resonances (Fig. 3). The incremental shifts indicate a fast exchange regime, on the NMR timescale, between the free and bound states. The dissociation constant (*K*_d_) was determined by fitting the CSPs as a function of diC_4_-PI(4,5)P_2_ concentration, which yielded *K*_d_ values of 7 and 306 μM at 0 and 0.1 M NaCl, respectively (Fig. S6 and Table 1). Interestingly, the affinity of diC_4_-PI(4,5)P_2_ to HTLV-1 myr(–)MA is ∼20-fold tighter than those observed for the HIV-1 and HIV-2 MA proteins (*K*_d_ ∼150 μM) (12,42), and > 120-fold tighter than that observed for the ASV MA protein (*K*_d_ = 850 μM) at 0 M NaCl. Among the signals that exhibited significant CSPs are Lys^47^, Lys^48^, and Lys^51^, which reside in helix II and form a basic patch on the surface of the protein (Fig. 3 and 4). Additionally, apparent CSPs were observed for several residues within helix II (Ser^39^, Ser^40^, Phe^43^, His^44^, Gln^45^, Leu^46^, Phe^49^, Leu^50^, and Ile^52^). Significant CSPs were also detected for signals corresponding to residues in the unstructured, N-terminal region of MA (Phe^5^, Ile^12^, Arg^14^, Arg^17^, Gly^18^, Leu^19^, Ala^20^, and Ala^21^; Fig. 4). Analysis of the surface electrostatic potential map of myr(–)MA_99_ shows that Arg^14^ and Arg^17^ (along with Arg^3^ and Arg^7^) form an extended second basic surface patch (Fig. 4).

**Table 1.** Dissociation constants for lipids and IPs binding to HTLV-1 myr(–)MA.

| **Lipid** | **$K_d$ (μM)** |
| --- | --- |
| diC<sub>4</sub>-PI(4,5)P<sub>2</sub> (0 M NaCl) | $7.2 \pm 2.1$ |
| diC<sub>4</sub>-PI(4,5)P<sub>2</sub> | $306 \pm 30$ |
| diC<sub>6</sub>-PI(4,5)P<sub>2</sub> | $195 \pm 19$ |
| I(1,4,5)P<sub>3</sub> | $338 \pm 42$ |
| I(1,4,5)P<sub>3</sub> (0 M NaCl) | $4.0 \pm 1.3$ |
| I(1,3,4,5)P<sub>4</sub> | $62 \pm 14$ |
| IP<sub>6</sub> | $23 \pm 11$ |
| IP<sub>6</sub> (0.2 M NaCl) | $36 \pm 7$ |
| IP<sub>6</sub> (0.4 M NaCl) | $295 \pm 56$ |

Titration were conducted in the presence of 0.1 M NaCl except when noted.

**Figure 3.**
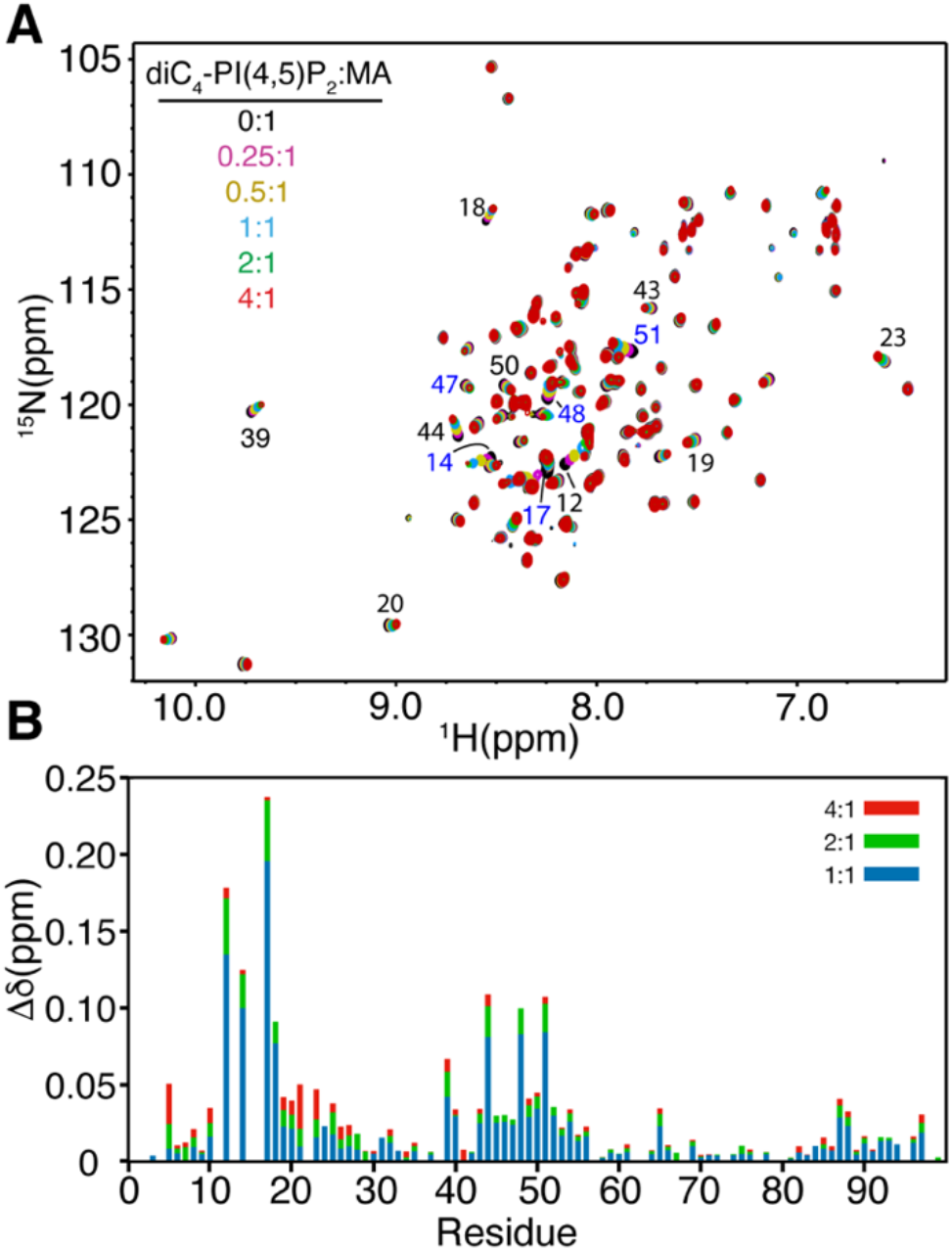
PI(4,5)P_2_ binding to myr(–)MA. *A*, Overlay of 2D ^1^H-^15^N HSQC spectra upon titration of myr(–)MA with diC_4_-PI(4,5)P_2_ [100 μM, 35 °C; diC_4_-PI(4,5)P_2_:MA = 0:1 (black), 0.25:1 (magenta), 0.5:1 (olive), 1:1 (cyan), 2:1 (green), 4:1 (red)] in 50 mM phosphates (pH 6) and 2 mM TCEP. *B*, Histogram of normalized ^1^H-^15^N chemical shift changes vs. residue number calculated from the HSQC spectra above.

**Figure 4.**
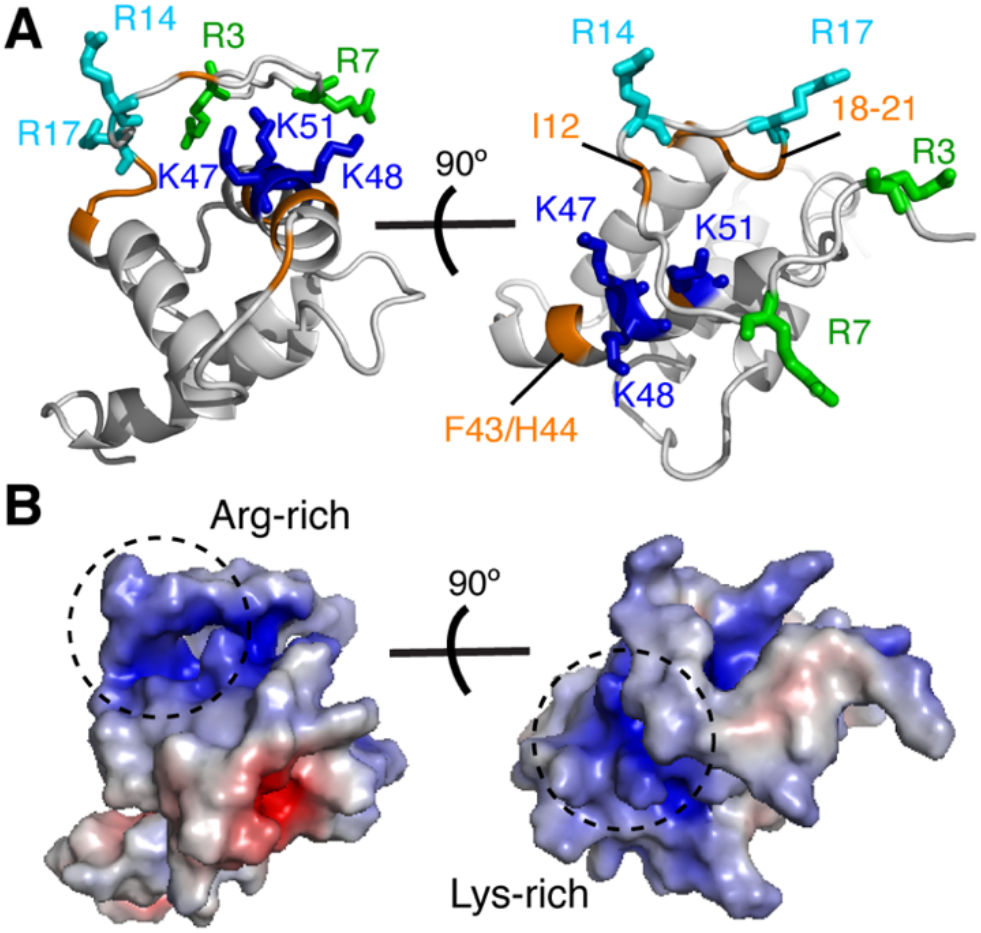
Chemical shift mapping of PI(4,5)P_2_ binding to myr(–)MA_99_. *A*, Cartoon representation of the myr(–)MA_99_ structure highlighting basic residues (blue and cyan) that exhibited substantial chemical shift changes upon binding of diC_4_-PI(4,5)P_2_. Signals of residues highlighted in orange are perturbed due to their proximity to the diC_4_-PI(4,5)P_2_ binding site. *B*, Electrostatic surface potential map of myr(–)MA_99_ showing the basic patches formed by the lysine-rich and arginine-rich regions.

Of note, several hydrophobic residues in the unstructured N-terminus (Phe^5^, Ile^12^, and Leu^19^) also exhibited CSPs upon titration of diC_4_-PI(4,5)P_2_. It is unclear whether these CSPs are a consequence of interactions with the polar head of diC_4_-PI(4,5)P_2_ via the N-terminal Arg^14^ and Arg^17^, or a result of direct contact with the acyl chains. It is possible that the flexibility of the N-terminus may allow for transient interactions with diC_4_-PI(4,5)P_2_ or other proximal regions. We assessed whether the acyl chains play a role in the interaction by conducting NMR titrations with dihexanoyl PI(4,5)P_2_ [diC_6_-PI(4,5)P_2_]. The CSPs for signals corresponding to hydrophobic residues in the N-terminus (Phe^5^, Ile^12^, and Leu^19^) and helix I (Phe^27^, Leu^28^, Ala^30^, Ala^31^, Tyr^32^) were slightly larger than those observed for diC_4_-PI(4,5)P_2_, suggesting that the acyl chains may interact with the N-terminal hydrophobic residues (Fig. S7). Consequently, a slightly higher binding affinity was observed for diC_6_-PI(4,5)P_2_ binding to myr(–)MA (Table 1). We do not rule out that the interactions between the acyl chains and the myr(–)MA protein are nonspecific.

To examine whether the polar head of PI(4,5)P_2_ is sufficient for myr(–)MA binding, we conducted NMR titrations with inositol 1,4,5-trisphosphate (IP_3_). As shown in Figure S8, ^1^H-^15^N resonances that exhibited significant CSPs are similar to those observed upon binding of diC_4_-PI(4,5)P_2_ (Fig. S7). IP_3_ titration data afforded a *K*_d_ of 4 μM, which is similar to that observed for diC_4_-PI(4,5)P_2_ (Table 1). Altogether, our data demonstrate that PI(4,5)P_2_ binds directly to HTLV-1 myr(–)MA via interactions between the polar head and a basic patch formed by lysine and arginine residues and that the acyl chains of PI(4,5)P_2_ are not critical for the interaction.

### Binding of myr(–)MA to IPs is governed by charge–charge interactions

To assess whether myr(–)MA binding to IPs is predominantly governed by electrostatic interactions, we conducted 2D HSQC NMR titration using IPs with varying number of phosphate groups on the inositol ring. Titration of myr(–)MA with inositol 1,3,4,5-tetrakisphosphate (IP_4_) yielded a *K*_d_ of 62 μM, which is ∼5-fold tighter than that observed for IP_3_ at the same buffer conditions (Table 1). Similar titration experiments performed with inositol hexakisphosphate (IP_6_) yielded a *K*_d_ of 23 μM, which again indicates that binding affinity is enhanced by increasing the number of phosphate groups on the inositol ring. Expectedly, increasing salt concentrations led to decrease in the binding affinity of IP_6_ to myr(–)MA (Table 1). Chemical shift mapping of the CSPs on the structure of myr(–)MA_99_ indicated that all tested PI(4,5)P_2_ and IPs bind to the same site. These results demonstrate that HTLV-1 myr(–)MA binding to PI(4,5)P_2_ is governed by ionic forces.

### Thermodynamics of IP binding to myr(–)MA

Having established the presence of a PI(4,5)P_2_ binding site on HTLV-1 MA, we sought to investigate the nature of the interaction and the contribution of enthalpic and entropic factors. To determine stoichiometry (*n*), enthalpy change (*ΔH°*) and entropic term (*TΔS°*), we conducted ITC experiments upon titration of myr(–)MA with IP_3_. We used IP_3_ because it is the polar headgroup of PI(4,5)P_2_ and because the binding affinity is sufficiently strong to yield analyzable ITC data. Applying a single set of identical sites model to fit the data (Fig. 5) yielded the following parameters: *K*_d_ = 3.7 ± 0.7 μM, *n* = 1.04 ± 0.03, Δ*H*° = -8.3 ± 0.3 kcal/mol, and *TΔS*° = 1.4 ± 0.1 kcal/mol. The ITC indicated that the *K*_d_ obtained by ITC data is very consistent with that obtained from the NMR titration data (Table 1), that myr(–)MA harbours a single IP binding site (*n ∼* 1), and that the exothermic reaction is indicative of the electrostatic nature of the interaction.

**Figure 5.**
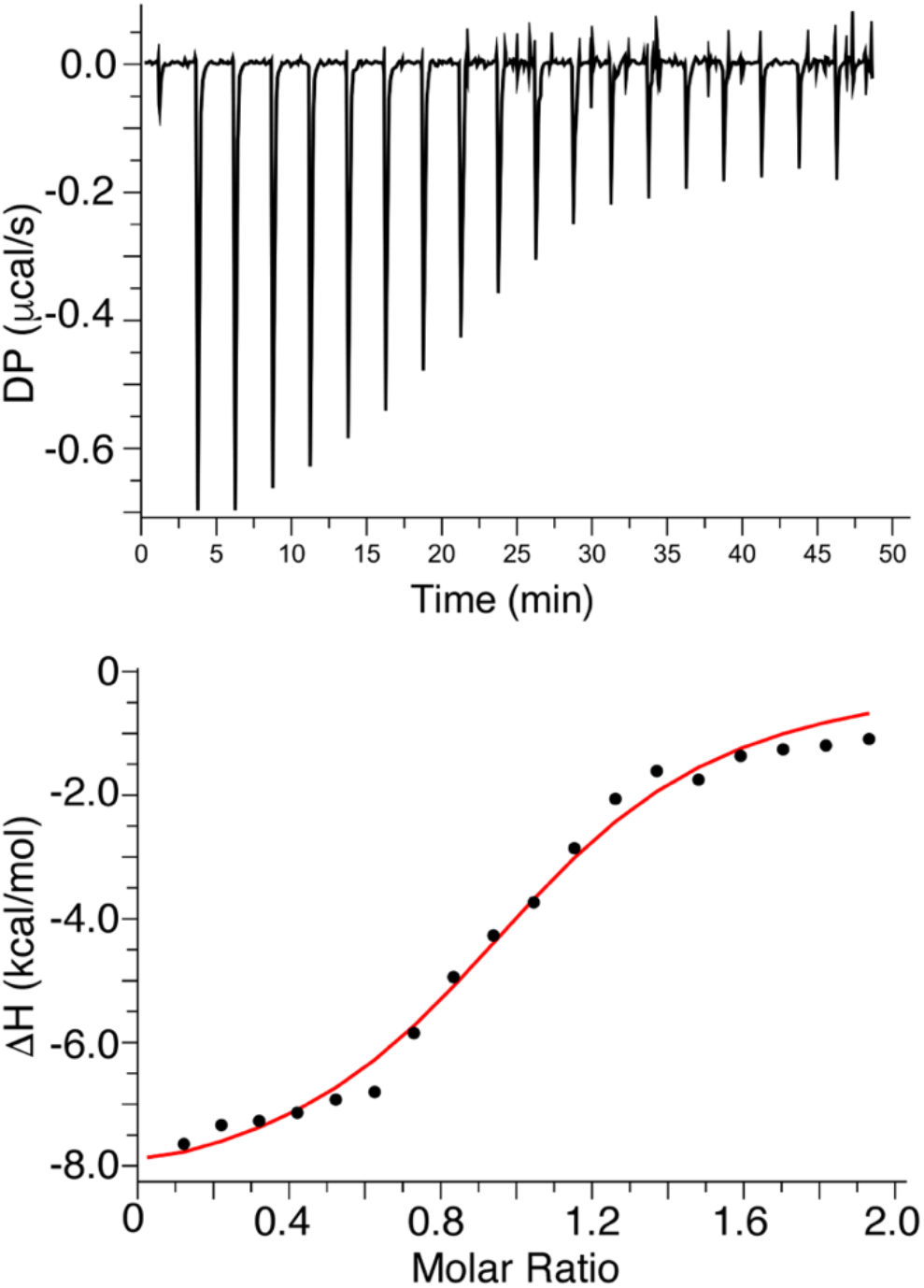
ITC data for binding of IP_3_ to myr(–)MA. ITC data obtained for titration of IP_3_ (400 μM) into myr(–)MA (40 μM) in 20 mM MES (pH 6.0) and 2 mM TCEP. Applying a single set of identical sites model to fit the data yielded a *K*_d_ of 3.7 ± 0.7 μM and stoichiometry value (*n*) of 1, indicating a single lipid binding site.

### Interaction of myr(–)MA with liposomes

The synergy between membrane components can influence the affinity of proteins to membranes. It is established that the affinity of retroviral MA and/or Gag proteins to membranes is enhanced by PS (6,19,22,26,62-67). Previous liposome binding data revealed that HTLV-1 Gag can bind PS-containing membranes efficiently even in the absence of PI(4,5)P_2_ (30). Herein, we employed a sensitive NMR–based assay to characterize binding of HTLV-1 myr(–)MA to large unilamellar vesicles (LUVs) containing native PI(4,5)P_2_ and PS. This NMR approach has been employed to characterize interaction of HIV-1 and ASV MA proteins to liposomes (40,47). The assay allows for measurement of the unbound protein population in solution under equilibrium conditions with the liposome–bound form (68,69). This assay can provide quantitative binding measurements such as *K*_d_ values and important information on the synergy of membrane components and cooperativity of binding. ^1^H NMR experiments were conducted on samples with fixed protein and total lipid concentrations, while varying lipid composition within LUVs. As predicted, myr(–)MA binding to LUVs made with 100% POPC was very weak, as judged by the negligible decrease in signal intensity (<10%) upon addition of saturating amounts of POPC LUVs (data not shown). However, NMR titrations of myr(–)MA with LUVs containing POPC and increasing amounts of PI(4,5)P_2_ led to increased signal attenuation (Fig. 6A), indicating that PI(4,5)P_2_ enhanced the affinity of myr(–)MA to liposomes. Data fitting using the Hill equation resulted in a *K*_d_ value of 36 μM (at 100 mM NaCl; Fig. 6B and Table 2), demonstrating that affinity of HTLV-1 myr(–)MA to membrane is enhanced by PI(4,5)P_2_ and in the absence of other acidic or charged lipids.

**Table 2.** Parameters for HTLV-1 myr(–)MA binding to LUVs determined using the Hill equation.

| **LUV** | **$K_d$ (μM)** | **n** |
| --- | --- | --- |
| POPC:POPS:PI(4,5)P<sub>2</sub> | 10 ± 1 | 0.8 |
| POPC:PI(4,5)P<sub>2</sub> | 36 ± 4 | 1.9 |
| POPC:diC<sub>16</sub>-PI(3,5)P<sub>2</sub> | 32 ± 2 | 1.7 |
| POPC:POPS | 118 ± 16 | 2.9 |

Titration were conducted in the presence of 0.1 M NaCl except when noted.

**Figure 6.**
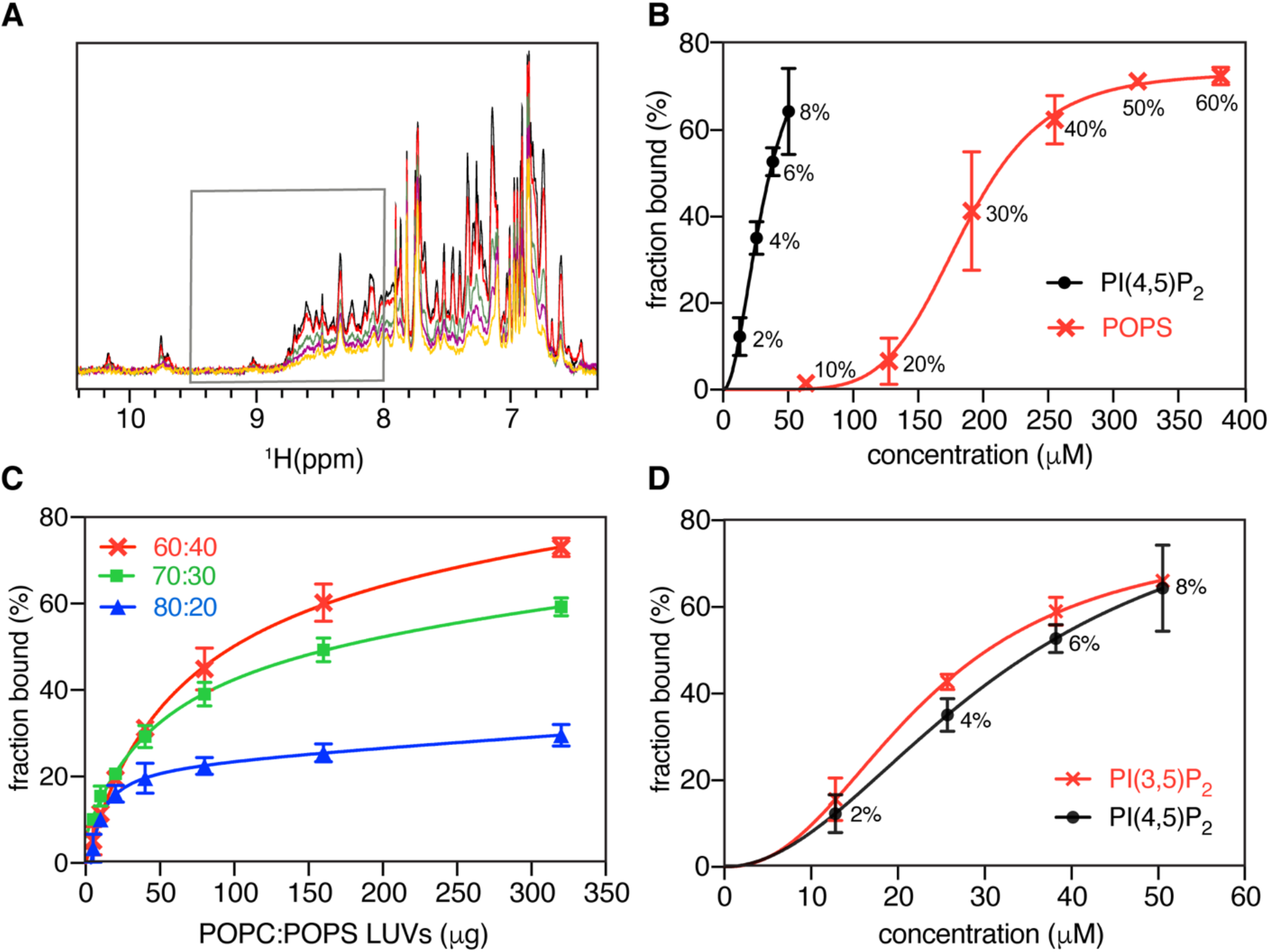
Interaction of myr(–)MA with LUVs. *A*, Overlay of ^1^H NMR spectra of myr(–)MA (50 µM) in the presence of fixed amount of LUVs with (100 – *x*)% mol. POPC and *x* = 0 (black), 2 (red), 4 (green), 6 (purple), and 8 (yellow) molar % PI(4,5)P_2_. Gray box marks the area of spectra integration. *B*, Isotherms of myr(–)MA binding to LUVs containing POPC and varying amounts of PI(4,5)P_2_ (black). Isotherms of myr(–)MA binding to LUVs containing POPC and varying %mol. of POPS is shown in red. Molar percentages of PI(4,5)P_2_ or POPS are indicated. Solid lines are Hill equation fits to the experimental data represented by points with error bars. *C*, myr(–)MA binding to POPC:POPS LUVs with varying concentrations of POPS as a function of increasing amounts of LUVs. *D*, Isotherms of myr(–)MA binding to PI(3,5)P_2_ and PI(4,5)P_2_ are shown in red and black respectively. Molar percentage of PI(4,5)P_2_ is indicated. Solid lines are fits of Hill equation to the experimental data represented by points with error bars.

Next, we examined the effect of PS on myr(–)MA binding by conducting NMR titrations with LUVs containing POPC and increasing amounts of POPS. Interestingly, in contrast to what was observed for other retroviral myr(–)MA proteins (40,47), binding was detectable upon incorporation of 20% mol. POPS (Figs. 6B and C). Fitting the binding isotherms with the Hill equation yielded a microscopic *K*_d_ of 118 μM for POPS (Table 2). Binding was highly cooperative as indicated by the *n* value (∼3) (Table 2), suggesting that myr(–)MA engages multiple PS molecules. Next, we conducted titrations with LUVs containing POPC, a fixed 20% mol. POPS and increasing amounts of PI(4,5)P_2_. Data fitting yielded a *K*_d_ of 10 μM, indicating a synergistic effect of PI(4,5)P_2_ and POPS in binding to myr(–)MA (Table 2).

### Specificity of myr(–)MA binding to phosphoinositides

To determine specificity of PI(4,5)P_2_ binding and whether the position of the phosphate groups on the inositol group affects myr(–)MA binding to membranes, titration experiments were conducted with POPC liposomes containing diC_16_-PI(3,5)P_2_, which differs from PI(4,5)P_2_ in the placement of a single phosphate. ^1^H NMR signals decreased in intensity upon addition of POPC/diC_16_-PI(3,5)P_2_ liposomes, similar to results obtained for myr(–)MA binding to POPC/PI(4,5)P_2_ liposomes. Fitting the binding data yielded a microscopic *K*_d_ of 32 μM, which is almost identical to that of PI(4,5)P_2_ (Fig. 6D and Table 2). These results indicate that myr(–)MA binding to LUVs is not dependent on the positioning of the phosphate group (4^th^ vs. 3^rd^ carbon) and that the overall charge of the lipid appears to be the major determinant of binding.

### Particle production and Gag subcellular localization

The studies above were conducted with the unmyristoylated MA protein. Previous studies demonstrated that PI(4,5)P_2_ binding to other retroviral MA proteins is independent of the myr group (12,40,42,61). It has also been shown that the overall binding affinity of HIV-1 MA to membranes is enhanced significantly (∼10-fold) by the myr group (19). However, recent NMR-based liposome studies have shown that HIV-1 myr(−)MA exhibited significant affinity for liposomes containing PI(4,5)P_2_, suggesting that the myr group contributes to initial membrane association but is not required for specific recognition of PIPs (47). We examined the role of the myr group in HTLV-1 Gag localization and particle production in mammalian cells by comparing the ability of wild-type (WT) and membrane binding deficient Gag (*i*.*e*., Gag protein containing the G2A mutation, which dramatically diminishes Gag binding to the plasma membrane) to produce immature particles. Immature VLPs were produced by transiently transfecting 293T cells with HTLV-1 Gag and Envelope (Env) expression plasmids, and particle production was confirmed by immunoblot. Particle production efficiency was analyzed for WT and G2A HTLV-1 Gag (Fig. 7A). Compared to WT Gag, the G2A Gag mutant which prevents Gag myristoylation, led to a 96% reduction of particle production compared to WT Gag (Fig. 7B). This observation is consistent with previous observations that a low level of HTLV-1 particle production is observed with Gag containing the G2A mutation (56), and that efficient membrane binding is requisite for robust particle production (70,71)

**Figure 7.**
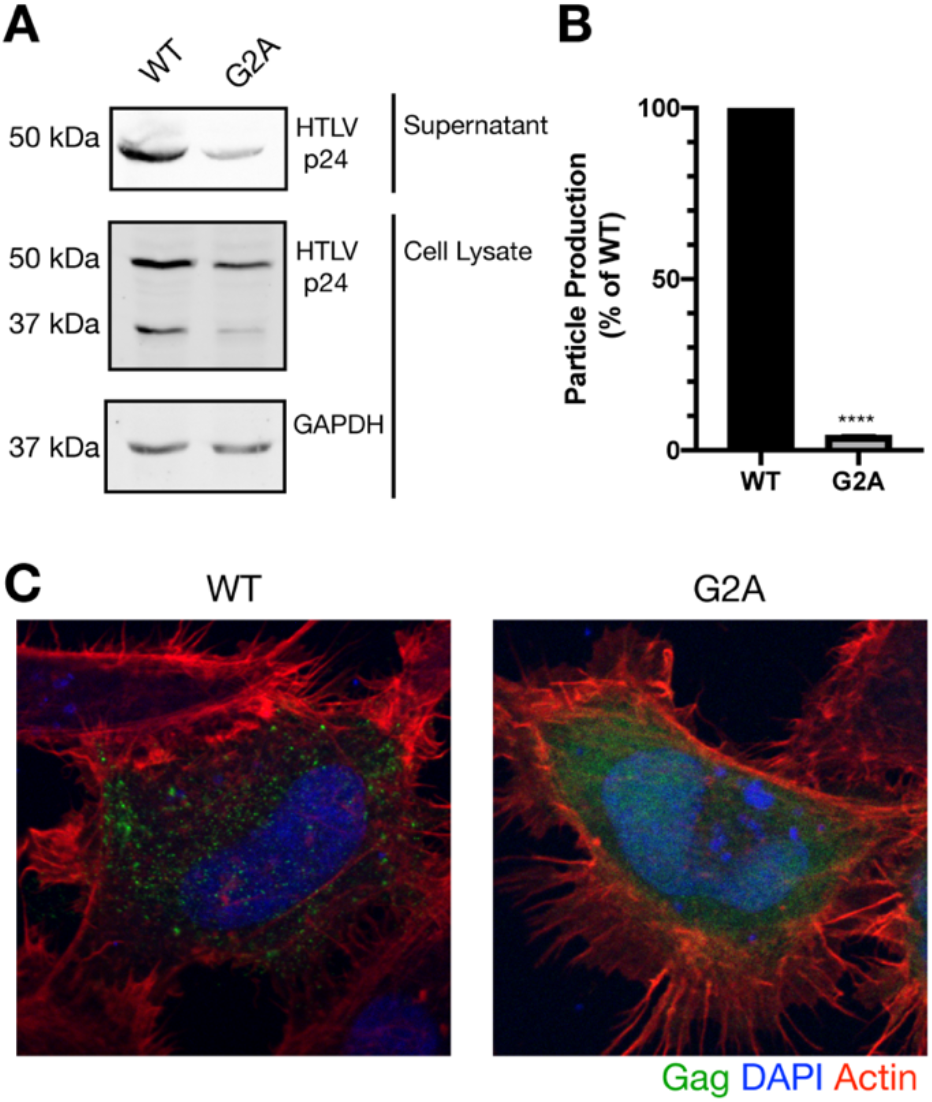
Analysis of particle production and HTLV-1 Gag subcellular distribution. *A*, Immunoblot analysis of particles released into the cell culture supernatant compared to Gag detected in lysates from particle-producing cells. Shown is a representative immunoblot from three independent biological replicates. Analysis of GAPDH in cell lysates was done to normalize samples. *B*, Quantification of particle production from three independent biological replicates. Relative particle production of the G2A Gag mutant is indicated as a percentage of WT Gag. **** indicates a p value ! 0.0001 (Student’s unpaired T-test). *C*, Gag subcellular distribution. Representative z-stack images from three independent biological replicates are shown in which Gag distribution in Hela cells was determined for the G2A Gag and WT Gag. Gag localization is indicated by green fluorescence; actin is indicated by red fluorescence; nuclei are indicated by blue (DAPI) fluorescence.

Next, confocal microscopy analysis was performed to investigate the role of the G2A mutation on Gag–membrane association compared to that of WT. Z-stacks of Hela cells expressing HTLV-1 WT Gag were found to exhibit punctate Gag fluorescence (Fig. 7C). The punctate Gag fluorescence, particularly at earlier time points post transfection, is indicative of Gag multimerization. In contrast, the diffuse pattern of fluorescence observed with the G2A Gag mutant was markedly distinct from that of WT (Fig. 7C). These observations indicate that WT Gag puncta are localized on the inner leaflet of the PM, while the G2A Gag mutant is diffuse and cytoplasmic.

## Discussion

Previous studies have shown that binding of HTLV-1 Gag to membranes is less dependent on PI(4,5)P_2_ than in HIV-1 (30). Unlike HIV-1 Gag, subcellular localization of HTLV-1 Gag and VLP release were minimally sensitive to overexpression of 5ptaseIV, suggesting that the interaction of HTLV-1 MA with PI(4,5)P_2_ is not essential for HTLV-1 particle assembly (30,31). On the other hand, it was found that inclusion of PI(4,5)P_2_ enhanced HTLV-1 Gag binding to liposomes but also bound efficiently to liposomes of similar negative charge upon substituting PI(4,5)P_2_ with PS (30). These results raised the questions whether the MA domain of HTLV-1 Gag contains a PI(4,5)P_2_ binding site, whether MA binds nonspecifically to acidic lipids including PI(4,5)P_2_, and/or whether Gag binding to membranes is mediated exclusively by charge-charge interactions. This study was designed to address these questions and to draw a comparison to previously studied retroviral MA proteins. To do so, we determined the structure of the membrane binding domain of MA [myr(–)MA_99_] and investigated the role of various factors driving membrane association such as the polar head and acyl chains of PI(4,5)P_2_, overall charge, lipid specificity, and membrane composition. Several important points have emerged from this study: (i) The structure of myr(–)MA_99_ revealed a PI(4,5)P_2_ binding site consisting of lysine and arginine residues. (ii) myr(–)MA binds to soluble analogs of PI(4,5)P_2_ with substantially higher affinity than previously studied retroviral MA proteins. (iii) The affinity of HTLV-1 myr(–)MA to liposomes is enhanced by PS and PI(4,5)P_2_, demonstrating the electrostatic nature of binding. (iv) The presence of PS in membranes enhances the binding affinity of myr(-)MA to PI(4,5)P_2_, indicating a synergistic effect between the two lipids. (v) HTLV-1 myr(–)MA has no preference to differentially phosphorylated forms of PIP_2_ since it bound to PI(3,5)P_2_ with a similar affinity to that of PI(4,5)P_2_. (vi) The acyl chains of PI(4,5)P_2_ have minimal role in the overall binding of PI(4,5)P_2_. (vii) Confocal microscopy data show that Gag is localized to the inner leaflet of the PM, while the Gag G2A mutant lacking myristoylation is diffuse and cytoplasmic. These findings provided significant insights into the mechanism by which the Gag protein binds to the inner leaflet of the PM and the subsequent production and release of HTLV-1 particles.

HTLV-1 Gag has been shown to bind to membranes with a higher affinity than that of HIV-1 Gag (30). Subsequent studies have shown that single amino acid substitutions that confer a large basic patch rendered HTLV-1 MA susceptible to the RNA-mediated block, suggesting that RNA blocks MA containing a large basic patch (31). These data supported a model in which HTLV-1 Gag localizes to the PM via the MA domain with higher efficiency but less specificity than for other retroviruses (30,31). It was proposed that Gag targeting is mediated by electrostatic interactions with acidic lipids with no specificity to PI(4,5)P_2_. Here, we show that all tested analogs of PI(4,5)P_2_ bind to the same site on myr(–)MA, which consists of Arg^14^, Arg^17^, Lys^47^, Lys^48^, and Lys^51^. Interestingly, the positioning of the three lysine residues on helix II of myr(–)MA is reminiscent of those observed for the ASV MA protein (40).

To our surprise, the affinity of HTLV-1 myr(–)MA to soluble analogs of PI(4,5)P_2_ is > 20-fold tighter than that observed for HIV-1 and HIV-2 MA, and ∼100-fold tighter than for ASV MA (Table 1). For the latter cases, Gag assembly has been shown to be dependent on PI(4,5)P_2_ (7,12,40). Tighter binding is likely attributed to the extensive H-bond capabilities of the guanidinium group of arginine (72), supporting a model in which Arg^14^ and Arg^17^ may play a significant role in enhancing PI(4,5)P_2_ binding.

The liposome binding assays, however, revealed that the affinity of HTLV-1 myr(–)MA to PI(4,5)P_2_ is similar to that observed for HIV-1 MA and ASV MA (40,47). It is likely that when not incorporated in membranes, PI(4,5)P_2_ analogs are better accommodated in the binding site on HTLV-1 MA. Additionally, a higher fraction of myr(–)MA bound to membranes containing PI(4,5)P_2_ vs. PS membranes of equivalent negative charge (Fig. 6B), indicating a slight preference of myr(–)MA to PI(4,5)P_2_. However, increasing PS ratio in liposomes resulted in similar levels of protein bound. Altogether, our studies confirmed the presence of PI(4,5)P_2_ binding site on HTLV-1 MA and demonstrated that binding is governed by electrostatic interactions.

Because previous studies demonstrated that PI(4,5)P_2_ binding to retroviral MA proteins is independent of the myr group (12,40,42,61), we examined the role of the myr group in HTLV-1 Gag colocalization on the PM and particle production in mammalian cells. The finding that G2A Gag mutant led to a 96% reduction of particle production compared to WT Gag (Fig. 7B) indicates that the myr group is required for proper Gag localization. This observation is consistent with previous observations that a low level of HTLV-1 particle production is observed with Gag containing the G2A mutation (56), and that efficient membrane binding is requisite for robust particle production (70,71). Confocal microscopy analysis of Hela cells expressing HTLV-1 WT Gag revealed formation of Gag puncta, particularly at earlier time points post transfection, indicative of Gag multimerization. In contrast, the G2A Gag mutant had a diffuse pattern of fluorescence (Fig. 7C). Altogether, these observations indicate that proper Gag multimerization and localization on the inner leaflet of the PM appear to be dependent on the myr group.

In a previous study, the effect of individual basic amino acid substitutions in the HTLV-1 MA protein on cell-to-cell transmission of the virus was examined. WT phenotype was only obtained for mutant viruses with mutations of Arg^7^ and Arg^97^ (73). However, point mutations of nine residues (Arg^3^, Arg^14^, Arg^17^, Arg^33^, Lys^47^, Lys^48^, Lys^51^, Lys^74^, and Arg^79^) completely abolished viral infectivity and impacted various steps of the replication cycle, including events following membrane targeting of Gag. Mapping of the basic residues on the structure of myr(–)MA_99_ revealed that the majority of the basic residues are located in the basic patch implicated in PI(4,5)P_2_ binding (Fig. S9). It was found that most of the mutations allowed normal synthesis, transport, and cleavage of the Gag precursor, but particle release was greatly affected for seven mutants (R3L, R14L, R17L, K48I, K74I, and R79L) (73). Interestingly, in situ immunofluorescence analysis of the distribution of the HTLV-1 Gag proteins in transfected cells revealed that the intracellular distributions of the Gag proteins with these point mutations were similar to that of the WT protein. Limited membrane binding studies using cell fractionation have shown that Gag R17L or K48I mutants bound to membranes with a similar affinity to the WT protein, suggesting that these two residues are important for infectivity at various stages of the viral replication cycle but do not play a major role, at least individually, in targeting the Gag precursor to the PM (73). Our structural data show that 5-7 basic residues can potentially contribute to membrane binding and that substitution of a single basic amino acid may not be detrimental to membrane binding.

A recent study demonstrated the capability of HIV-1 MA to partially displace the acyl chains from the bilayer membrane via interaction with hydrophobic regions of MA and stabilization of the MA-lattice ((48)). This is consistent with the increasing binding affinities of HIV-1 MA to truncated forms of PI(4,5)P_2_ with increasing acyl chain length ((42). In this study, we observed that the binding affinity of PI(4,5)P_2_ to HTLV-1 myr(–)MA does not depend on the presence/lack of or length of acyl chain (Table 1). A modest increase in affinity (∼2-fold) is observed upon increasing the length of the acyl chain by two methylene groups. As revealed by the CSPs (Fig. S7), this slight increase in the affinity is probably caused by interactions between the acyl chains and hydrophobic residues in the unstructured N-terminal region and helix I of myr(–)MA. Further studies are needed to investigate whether the interaction of MA with the acyl chain can occur in infected cells and can overcome the energy penalty resulting from displacement of part of the acyl chain from the membrane bilayer, as has been shown for HIV-1 (48).

Our data show no preferential binding of PI(4,5)P_2_ to myr(–)MA compared to PI(3,5)P_2_, indicating that the position of the phosphate groups is not a key determinant for binding to HTLV-1 myr(–)MA. Interestingly, this result is similar to that obtained for HIV-1 and ASV MA in which the affinity to PI(3,5)P_2_ was found to be relatively similar to PI(4,5)P_2_ (40,47). It has been suggested that since PI(3,5)P_2_ abundance in cells is significantly lower than that of PI(4,5)P_2_ (∼100-fold lower (74)), PM targeting of Gag results from the high relative concentration of PI(4,5)P_2_ rather than differences in affinity of MA for these phosphoinositides (47). This hypothesis is perhaps applicable to HTLV-1 Gag since it appears that the total negative charge on lipids is more important than the positioning of the phosphate groups.

In summary, our data support a model in which HTLV-1 MA binding to membranes is governed by electrostatic interactions and that the affinity of MA binding to membranes is enhanced by acidic lipids such as PS and PI(4,5)P_2_. These findings provide a structural framework for HTLV-1 Gag binding to the inner leaflet of the PM and advance our understanding of the basic mechanisms of retroviral assembly.

## Experimental procedures

### Sample Preparation

*Plasmid construction*. The *MA* genes encoding for amino acids 1-130 or 1-99 were generated via PCR using a plasmid containing the full-length HTLV-1 *Gag* gene as a template (75). *MA* genes were inserted into a pET11a vector using standard cloning techniques, yielding a construct that is fused to a *His*_*6*_*-tag* gene on the 3’-end. Plasmid sequencing was performed at the Heflin Genomics Core at the University of Alabama at Birmingham.

*Protein expression and purification*. The myr(–)MA and myr(–)MA_99_ proteins were overexpressed in *Escherichia coli* BL21 Codon Plus-RIL cells (Agilent Technologies). Cells were grown in LB broth supplemented with 100 mg/L Ampicillin at 37 °C. Cells were induced with isopropyl β-D-1-thiogalactoside when the OD_600_ was ∼0.7 and grown at 22 °C overnight. Cells were harvested via centrifugation and stored at –80 °C. Cell pellet was resuspended in lysis buffer (50 mM sodium phosphates, pH 8.0, 500 mM NaCl, 40 Mm imidazole, 1% Triton, and 1 mM phenylmethylsulfonyl fluoride (PMSF)). Cells were lysed via sonication and lysate was spun down at 35,000xg for 30 min. The supernatant was subjected to cobalt affinity chromatography and eluted with imidazole gradient. Proteins were further purified by cation-exchange and size-exclusion chromatography. Protein samples were dialyzed in NMR buffer (20 mM MES, pH 6.0, 100 mM NaCl, and 2 mM TCEP) or LUV buffer (50 mM sodium phosphates, pH 7.4, and 100 mM NaCl) and concentrated using 3 kDa cut-off centrifugal filter units. Uniformly ^15^N and ^15^N-,^13^C-labeled MA samples were prepared by growing cells in M9 minimal medium containing ^15^NH_4_Cl and glucose-^13^C_6_. Protein purification was performed as described above.

*Gel filtration assay*. The mobility of myr(–)MA and myr(–)MA_99_ proteins was analyzed by a gel filtration assay. Briefly, 0.5 mL of 100 μM protein samples was loaded on ENrich SEC 70 column (BioRad) in a buffer containing 50 mM phosphates (pH 7.4), 300 mM NaCl and 2mM TCEP. Protein fractions were analyzed by SDS-PAGE and stained by Coomassie brilliant blue. The approximate molecular weights of the loaded proteins were determined by molecular weight calibration kits (GE Healthcare).

### Preparation of Large Unilamellar Vesicles (LUVs)

1-Palmitoyl-2-oleoyl-*sn*-glycero-3-phosphocholine (POPC), 1-palmitoyl-2-oleoyl-*sn*-glycero-3-phospho-L-serine (POPS), porcine brain PI(4,5)P_2_ (Avanti Polar Lipids), and dipalmitoyl-phosphatidylinositol 3,5-bisphosphate [diC_16_-PI(3,5)P_2_] (Echelon Biosciences) were used as received. Lipids were mixed in appropriate ratios and solvent was evaporated under a stream of air, followed by lyophilization. Dried lipids were then resuspended in a buffer containing 50 mM sodium phosphates (pH 7.4) and 100 mM NaCl by repeated brief vortexing and allowed to rehydrate for 45 min at room temperature. Lipid suspension was then passed 30 times through a 100 nm pore filter in an extruder (Avanti Polar Lipids). LUV’s were stored at 4 °C and used within 24h. Final total lipid concentration in LUV stocks was 10 mg/ml.

### NMR Spectroscopy

NMR data were collected at 35 °C on a Bruker Avance II (700 MHz ^1^H) equipped with a cryogenic triple-resonance probe, processed with NMRPIPE (76) and analyzed with NMRVIEW (77) or CCPN Analysis (78). ^13^C-, ^15^N-, or ^13^C-/^15^N-labeled protein samples were prepared at ∼ 300-500 μM in 50 mM sodium phosphates (pH 6.0), 100 mM NaCl, and 1 mM TCEP. ^1^H, ^13^C and ^15^N resonances were assigned using ^1^H–^15^N HSQC, HNCA, HN(CO)CA, HNCACB, HN(CO)CACB, HNCO, ^15^N-edited HSQC-TOCSY, ^15^N-edited HSQC-NOESY and (H)CCH-TOCSY experiments. ^15^N-edited NOESY-HSQC and ^13^C-edited HMQC-NOESY data were collected with a mixing time of 120 ms.

### Lipid NMR titrations

^1^H-^15^N HSQC NMR titrations were conducted with 50-100 μM samples of ^15^N-labeled myr(–)MA_99_ or myr(–)MA in 20 mM MES (pH 6.0), 2 mM TCEP, and varying NaCl concentrations. Stock solutions of lipids were prepared in water at 10-50 mM. The pH of IP_6_ stock solution was adjusted to 6.0 by using NaOH prior to titrations. CSPs were calculated as 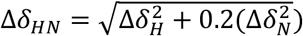, where Δδ_*H*_ and Δδ_*N*_ are ^1^H and ^15^N chemical shift changes, respectively. Dissociation constants were calculated by non-linear least-square fitting algorithm in gnuplot software (http://www.gnuplot.info) using the equation:

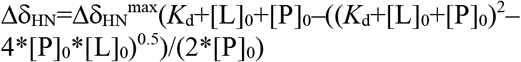

where 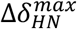 is chemical shift difference between complex and free protein, [L]_0_ total concentration of lipid, and [P]_0_ total concentration of protein.

### LUV NMR titration

Individually prepared samples for NMR titration contained 25 or 50 μM myr(–)MA in 50 mM sodium phosphates (pH 7.4), 100 mM NaCl, 2 mM TCEP, 250 or 500 μg LUVs with varying POPS and/or PI(4,5)P_2_ concentrations, and 5% D_2_O (vol/vol) in a total volume of 500 μL. ^1^H NMR spectra with excitation sculpting water suppression were recorded for each sample and integral intensity measured in the region 9.5–8.0 ppm. The amount of protein bound to LUVs was determined as the difference between integrals of samples with and without LUVs. The binding data (averages of 2-3 experiments) were fitted in Matlab 2015b (MathWorks) using Hill equation 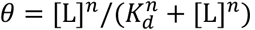, where *θ* is fraction of bound protein, [L] concentration of available lipid in LUV, *K*_*d*_ microscopic dissociation constant, and *n* cooperativity constant.

### Isothermal Titration Calorimetry

Thermodynamic parameters of IP_3_ binding to myr(–)MA were determined using a MicroCal PEAQ-ITC (Malvern Instruments). ITC experiments were conducted in a buffer containing 20 mM MES (pH 6.0) and 2 mM TCEP. IP_3_ prepared at 400 μM in the same buffer was titrated into 40 μM myr(–)MA. Heat of reaction was measured at 25 °C for 19 injections. Heat of dilution was measured by titrating IP_3_ into buffer and was subtracted from the heat of binding. Data analysis was performed using PEAQ analysis software. The thermodynamic parameters were determined by fitting baseline-corrected data by a binding model for a single set of identical sites.

### Structure Calculations

Structure calculations were performed using Unio’10 software (79) that utilizes Atnos/Candid functionality for automated iterative peak picking of raw NOESY spectra, peak assignments and calibration, in conjunction with CYANA structure calculation engine (80,81). Backbone φ and Ψ dihedral angle constraints were generated by Unio based on the chemical shifts. Tolerance windows for direct and indirect dimensions were set to 0.04. NOESY spectra were converted to XEASY format using CARA (82). Structures were visualized in Pymol (Schrödinger, LLC.). Electrostatic potential maps were generated using PDB2PQR and APBS software complied within Pymol (83,84).

### Particle production assay

HEK 293T cells (NIH HIV Reagent Program) and Hela cells (ATCC, VA) were grown in Dulbecco’s-modified Eagles medium (Corning, NY) supplemented with 10% fetal clone III (Cytiva Life Sciences, MA). For cell culture assays, a codon-optimized HTLV-1 Gag (pN3 HTLV-1 Gag and pN3 HTLV-1 Gag-EYFP) (75), or a derivative containing the G2A mutation (pN3 HTLV-1 G2A Gag and pN3 HTLV-1 G2A Gag-EYFP) (56) were used. The HTLV-1 Env expression construct was graciously provided by Kathryn Jones and Marie-Christine Dokhelar (85).

HEK 293T cells were co-transfected with 8 µg pN3 HTLV-1 Gag and 0.8 µg HTLV-1 Env plasmids by using the polyethylenimine (PEI) transfection reagent (Sigma-Aldrich, MO) at a 1:3 weight to volume ratio. Forty-eight hours post-transfection, the cell culture supernatant was filtered with a 0.22 µm filter and concentrated for 90 minutes by ultracentrifugation using a 50.2 Ti rotor at 150,000 x *g* through a 5 mL 8% OptiPrep (Sigma-Aldrich, MO) cushion to collect virus-like particles (VLPs). VLPs were resuspended in PBS + 0.1% Triton X-100 and concentrated 100x the initial volume. Cells were harvested, washed with PBS and lysed in PBS + 0.1% Triton X-100. Cell lysates and cell culture supernatants were normalized for protein concentration using the BCA assay according to the manufacturers’ instructions (Thermo Fisher Scientific, MA). Each sample (30 µg) was run on a 14% SDS-PAGE gel and transferred to a nitrocellulose membrane (Bio-Rad, CA). HTLV-1 Gag expression in cells and VLPs in the cell culture supernatant was detected using a mouse monoclonal antibody HTLV-1 p24 (6G9) (Santa Cruz Biotechnology, TX) 1:3,000 and secondary antibody goat anti-mouse IgG StarBright Blue 700 (Bio-Rad) 1:2500. GAPDH in cell lysates was detected using an anti-GAPDH hFAB Rhodamine antibody (Bio-Rad) 1:3000. Immunoblots were imaged by using a ChemiDoc Touch system (Bio-Rad) and analyzed with ImageLab (Bio-Rad).

### Confocal microscopy analysis of Gag subcellular distribution

Hela cells were co-transfected with pN3 HTLV-1 Gag, pN3 HTLV-1 Gag-EYFP, and pHTLV-1 Env at a 3:1:0.4 ratio using PEI at a 1:3 weight to volume ratio. Sixteen hours post-transfection, cells were fixed with 4% paraformaldehyde and permeabilized. The cytoskeleton was labeled with ActinRed 555 ReadyProbes Reagent and cell nuclei were stained with NucBlue Fixed Cell ReadyProbes Reagent (Life Technologies, CA) according to the manufacturers’ instructions. Cells were imaged using a Zeiss LSM700 confocal laser scanning microscope with a Plan-Apochromat 63x/1.4 aperture (NA) oil objective.

## Data Availability

The atomic coordinates of HTLV-1 myr(–)MA_99_ (code 7M1W) have been deposited in the Protein Data Bank (http://wwpdb.org/).

*The NMR chemical shift data* for *HTLV-1 myr(–)MA are available from the Biological Magnetic Resonance Data Bank under BMRB accession number 30880*.

The raw data described in the manuscript can be shared upon request by directly contacting.

## Supporting information

This article contains supporting information.

## Acknowledgments

We thank the O’Neal Comprehensive Cancer Center at the University of Alabama at Birmingham (funded by the NCI grant P30 CA013148) for supporting the High-Field NMR facility.

## Author contributions

DH, LWZ, HMH, NAW, LMM, and JSS designed the experiments. DH and LWZ expressed, purified, and characterized the proteins. DH and LWZ performed the NMR and ITC experiments and analyzed the results. HMH, NAW, and LMM performed the confocal microscopy data and virus production assays. DH, HMH, NAW, LMM and JSS wrote the paper. DH, LWZ, HMH, NAW, LMM, and JSS edited the paper.

## Abbreviations

HTLV-1: human T-cell leukemia virus type 1
MA: myristoylated matrix
myr(–)MA: unmyristoylated matrix
PI(4,5)P_2_: phosphatidylinositol 4,5-bisphosphate
NMR: nuclear magnetic resonance
HSQC: heteronuclear single quantum coherence
CSP: chemical shift perturbation
ITC: isothermal titration calorimetry
PI(4,5)P_2_: phosphatidylinositol 4,5-bisphosphate
PI(3,5)P_2_: phosphatidylinositol 3,5-bisphosphate
IP_3_: inositol 1,4,5-trisphosphate
IP_4_: inositol 1,3,4,5-tetrakisphosphate
IP_6_: inositol hexakisphosphate
POPC: 1-palmitoyl-2-oleoyl-sn-glycero-3-phosphocholine
POPS: 1-palmitoyl-2-oleoyl-sn-glycero-3-phospho-L-serine
IP: inositol phosphate
LUV: large unilamellar vesicle.

## References

1. Finzi, A., Orthwein, A., Mercier, J., and Cohen, E. A. (2007) Productive Human Immunodeficiency Virus Type 1 Assembly Takes Place at the Plasma Membrane. J. Virol. 81, 7476–7490

2. Gousset, K., Ablan, S. D., Coren, L. V., Ono, A., Soheilian, F., Nagashima, K., Ott, D. E., and Freed, E. O. (2008) Real-time visualization of HIV-1 GAG trafficking in infected macrophages. PLoS Pathog. 4, e1000015

3. Joshi, A., Ablan, S. D., Soheilian, F., Nagashima, K., and Freed, E. O. (2009) Evidence that productive human immunodeficiency virus type 1 assembly can occur in an intracellular compartment J. Virol. 83, 5375–5387

4. Jouvenet, N., Neil, S. J. D., Bess, C., Johnson, M. C., Virgen, C. A., Simon, S. M., and Bieniasz, P. D. (2006) Plasma membrane is the site of productive HIV-1 particle assembly. PLoS Biol. 4, e435

5. Welsch, S., Keppler, O. T., Habermann, A., Allespach, I., Krijnse-Locker, J., and Kräusslich, H.-G. (2007) HIV-1 buds predominantly at the plasma membrane of primary human macrophages. PLoS Pathog. 3, e36

6. Chukkapalli, V., Hogue, I. B., Boyko, V., Hu, W.-S., and Ono, A. (2008) Interaction between HIV-1 Gag matrix domain and phosphatidylinositol-(4,5)-bisphosphate is essential for efficient Gag-membrane binding. J. Virol. 82, 2405–2417

7. Ono, A., Ablan, S. D., Lockett, S. J., Nagashima, K., and Freed, E. O. (2004) Phosphatidylinositol (4,5) bisphosphate regulates HIV-1 Gag targeting to the plasma membrane. Proc Natl Acad Sci U S A 101, 14889–14894

8. Ghanam, R. H., Samal, A. B., Fernandez, T. F., and Saad, J. S. (2012) Role of the HIV-1 matrix protein in Gag intracellular trafficking and targeting to the plasma membrane for virus assembly. Front Microbiol. 3, 55

9. Vlach, J., and Saad, J. S. (2015) Structural and molecular determinants of HIV-1 Gag binding to the plasma membrane. Front Microbiol 6, 232

10. Hamard-Peron, E., Juillard, F., Saad, J. S., Roy, C., Roingeard, P., Summers, M. F., Darlix, J. L., Picart, C., and Muriaux, D. (2010) Targeting of murine leukemia virus gag to the plasma membrane is mediated by PI(4,5)P2/PS and a polybasic region in the matrix. J Virol 84, 503–515

11. Prchal, J., Kroupa, T., Ruml, T., and Hrabal, R. (2014) Interaction of Mason-Pfizer monkey virus matrix protein with plasma membrane. Front Microbiol 4, 423

12. Saad, J. S., Ablan, S. D., Ghanam, R. H., Kim, A., Andrews, K., Nagashima, K., Soheilian, F., Freed, E. O., and Summers, M. F. (2008) Structure of the myristylated HIV-2 MA protein and the role of phosphatidylinositol-(4,5)-bisphosphate in membrane targeting. J. Mol. Biol. 382, 434–447

13. Freed, E. O. (2015) HIV-1 assembly, release and maturation. Nat Rev Microbiol 13, 484–496

14. Ganser-Pornillos, B. K., Yeager, M., and Sundquist, W. I. (2008) The structural biology of HIV assembly. Curr Opin Struct Biol 18, 203–217

15. Chukkapalli, V., Inlora, J., Todd, G. C., and Ono, A. (2013) Evidence in support of RNA-mediated inhibition of phosphatidylserine-dependent HIV-1 Gag membrane binding in cells. J. Virol. 87, 7155–7159

16. Chukkapalli, V., and Ono, A. (2011) Molecular Determinants that Regulate Plasma Membrane Association of HIV-1 Gag. J. Mol. Biol. 410, 512–524

17. Purohit, P., Dupont, S., Stevenson, M., and Green, M. R. (2001) Sequence-specific interaction between HIV-1 matrix protein and viral genomic RNA revealed by in vitro genetic selection. RNA 7, 576–584

18. Li, H., Dou, J., Ding, L., and Spearman, P. (2007) Myristoylation is required for human immunodeficiency virus type 1 Gag-Gag multimerization in mammalian cells. J. Virol. 81, 12899–12910

19. Dalton, A. K., Ako-Adjei, D., Murray, P. S., Murray, D., and Vogt, M. V. (2007) Electrostatic Interactions Drive Membrane Association of the Human Immunodeficiency Virus Type 1 Gag MA Domain. J. Virol. 81, 6434–6445

20. Dick, R. A., Goh, S. L., Feigenson, G. W., and Vogt, V. M. (2012) HIV-1 Gag protein can sense the cholesterol and acyl chain environment in model membranes. Proc. Natl. Acad. Sci. U.S.A. 109, 18761–18767

21. Waheed, A. A., and Freed, E. O. (2009) Lipids and membrane microdomains in HIV-1 replication. Virus Res. 143, 162–176

22. Chukkapalli, V., Oh, S. J., and Ono, A. (2010) Opposing mechanisms involving RNA and lipids regulate HIV-1 Gag membrane binding through the highly basic region of the matrix domain. Proc. Natl. Acad. Sci. 107, 1600–1605

23. Waheed, A. A., and Freed, E. O. (2010) The role of lipids in retrovirus replication. Viruses 2, 1146–1180

24. Ono, A. (2010) HIV-1 assembly at the plasma membrane. Vaccine 28 Suppl 2, B55–59

25. Chan, J., Dick, R. A., and Vogt, V. M. (2011) Rous Sarcoma Virus Gag Has No Specific Requirement for Phosphatidylinositol-(4,5)-Bisphosphate for Plasma Membrane Association In Vivo or for Liposome Interaction In Vitro. J. Virol. 85, 10851–10860

26. Alfadhli, A., Still, A., and Barklis, E. (2009) Analysis of human immunodeficiency virus type 1 matrix binding to membranes and nucleic acids. J Virol 83, 12196–12203

27. Barros, M., Heinrich, F., Datta, S. A., Rein, A., Karageorgos, I., Nanda, H., and Losche, M. (2016) Membrane Binding of HIV-1 Matrix Protein: Dependence on Bilayer Composition and Protein Lipidation. J Virol 90, 4544–4555

28. Olety, B., Veatch, S. L., and Ono, A. (2015) Phosphatidylinositol-(4,5)-Bisphosphate Acyl Chains Differentiate Membrane Binding of HIV-1 Gag from That of the Phospholipase Cdelta1 Pleckstrin Homology Domain. J Virol 89, 7861–7873

29. Gaines, C. R., Tkacik, E., Rivera-Oven, A., Somani, P., Achimovich, A., Alabi, T., Zhu, A., Getachew, N., Yang, A. L., McDonough, M., Hawkins, T., Spadaro, Z., and Summers, M. F. (2018) HIV-1 Matrix Protein Interactions with tRNA: Implications for Membrane Targeting. J Mol Biol 430, 2113–2127

30. Inlora, J., Chukkapalli, V., Derse, D., and Ono, A. (2011) Gag localization and virus-like particle release mediated by the matrix domain of human T-lymphotropic virus type 1 Gag are less dependent on phosphatidylinositol-(4,5)-bisphosphate than those mediated by the matrix domain of HIV-1 Gag. J Virol 85, 3802–3810

31. Inlora, J., Collins, D. R., Trubin, M. E., Chung, J. Y., and Ono, A. (2014) Membrane binding and subcellular localization of retroviral Gag proteins are differentially regulated by MA interactions with phosphatidylinositol-(4,5)-bisphosphate and RNA. mBio 5, e02202

32. Thornhill, D., Olety, B., and Ono, A. (2019) Relationships between MA-RNA binding in cells and suppression of HIV-1 Gag mislocalization to intracellular membranes. J Virol

33. Thornhill, D., Murakami, T., and Ono, A. (2020) Rendezvous at Plasma Membrane: Cellular Lipids and tRNA Set up Sites of HIV-1 Particle Assembly and Incorporation of Host Transmembrane Proteins. Viruses 12

34. Hammond, G. R., Fischer, M. J., Anderson, K. E., Holdich, J., Koteci, A., Balla, T., and Irvine, R. F. (2012) PI4P and PI(4,5)P2 are essential but independent lipid determinants of membrane identity. Science 337, 727–730

35. Kolay, S., Basu, U., and Raghu, P. (2016) Control of diverse subcellular processes by a single multi-functional lipid phosphatidylinositol 4,5-bisphosphate [PI(4,5)P2]. Biochem J 473, 1681–1692

36. Stansell, E., Apkarian, R., Haubova, S., Diehl, W. E., Tytler, E. M., and Hunter, E. (2007) Basic residues in the Mason-Pfizer monkey virus gag matrix domain regulate intracellular trafficking and capsid-membrane interactions. J. Virol. 81, 8977–8988

37. Prchal, J., Srb, P., Hunter, E., Ruml, T., and Hrabal, R. (2012) The Structure of Myristoylated Mason-Pfizer Monkey Virus Matrix Protein and the Role of Phosphatidylinositol-(4,5)-Bisphosphate in Its Membrane Binding. J. Mol. Biol. 423, 427–438

38. Brown, L. A., Cox, C., Baptiste, J., Summers, H., Button, R., Bahlow, K., Spurrier, V., Kyser, J., Luttge, B. G., Kuo, L., Freed, E. O., and Summers, M. F. (2015) NMR structure of the myristylated feline immunodeficiency virus matrix protein. Viruses 7, 2210–2229

39. Nadaraia-Hoke, S., Bann, D. V., Lochmann, T. L., Gudleski-O’Regan, N., and Parent, L. J. (2013) Alterations in the MA and NC domains modulate phosphoinositide-dependent plasma membrane localization of the Rous sarcoma virus Gag protein. J. Virol. 87, 3609–3615

40. Vlach, J., Eastep, G. N., Ghanam, R. H., Watanabe, S. M., Carter, C. A., and Saad, J. S. (2018) Structural basis for targeting avian sarcoma virus Gag polyprotein to the plasma membrane for virus assembly. J Biol Chem 293, 18828–18840

41. Watanabe, S. M., Medina, G. N., Eastep, G. N., Ghanam, R. H., Vlach, J., Saad, J. S., and Carter, C. A. (2018) The matrix domain of the Gag protein from avian sarcoma virus contains a PI(4,5)P2-binding site that targets Gag to the cell periphery. J Biol Chem 293, 18841–18853

42. Saad, J. S., Miller, J., Tai, J., Kim, A., Ghanam, R. H., and Summers, M. F. (2006) Structural basis for targeting HIV-1 Gag proteins to the plasma membrane for virus assembly. Proc Natl Acad Sci U S A 103, 11364–11369

43. Vlach, J., and Saad, J. S. (2013) Trio engagement via plasma membrane phospholipids and the myristoyl moiety governs HIV-1 matrix binding to bilayers. Proc Natl Acad Sci U S A 110, 3525–3530

44. Murphy, R. E., Samal, A. B., Vlach, J., Mas, V., Prevelige, P. E., and Saad, J. S. (2019) Structural and biophysical characterizations of HIV-1 matrix trimer binding to lipid nanodiscs shed light on virus assembly. J Biol Chem 294, 18600–18612

45. Anraku, K., Fukuda, R., Takamune, N., Misumi, S., Okamoto, Y., Otsuka, M., and Fujita, M. (2010) Highly sensitive analysis of the interaction between HIV-1 Gag and phosphoinositide derivatives based on surface plasmon resonance. Biochemistry 49, 5109–5116

46. Charlier, L., Louet, M., Chaloin, L., Fuchs, P., Martinez, J., Muriaux, D., Favard, C., and Floquet, N. (2014) Coarse-Grained Simulations of the HIV-1 Matrix Protein Anchoring: Revisiting Its Assembly on Membrane Domains. Biophys. J. 106, 577–585

47. Mercredi, P. Y., Bucca, N., Loeliger, B., Gaines, C. R., Mehta, M., Bhargava, P., Tedbury, P. R., Charlier, L., Floquet, N., Muriaux, D., Favard, C., Sanders, C. R., Freed, E. O., Marchant, J., and Summers, M. F. (2016) Structural and Molecular Determinants of Membrane Binding by the HIV-1 Matrix Protein. J Mol Biol 428, 1637–1655

48. Qu, K., Ke, Z., Zila, V., Anders-Össwein, M., Glass, B., Müller, B., Kräusslich, H.-G., and Briggs, J. A. G. (2021) Maturation of the matrix and viral membrane of HIV-1. bioRxiv, 10.1101/2020.1109.1123.309542

49. Martinez, M. P., Al-Saleem, J., and Green, P. L. (2019) Comparative virology of HTLV-1 and HTLV-2. Retrovirology 16, 21

50. Poiesz, B. J., Ruscetti, F. W., Reitz, M. S., Kalyanaraman, V. S., and Gallo, R. C. (1981) Isolation of a new type C retrovirus (HTLV) in primary uncultured cells of a patient with Sezary T-cell leukaemia. Nature 294, 268–271

51. Yoshida, M., Miyoshi, I., and Hinuma, Y. (1982) Isolation and characterization of retrovirus from cell lines of human adult T-cell leukemia and its implication in the disease. Proc Natl Acad Sci U S A 79, 2031–2035

52. Ceccaldi, P. E., Delebecque, F., Prevost, M. C., Moris, A., Abastado, J. P., Gessain, A., Schwartz, O., and Ozden, S. (2006) DC-SIGN facilitates fusion of dendritic cells with human T-cell leukemia virus type 1-infected cells. J Virol 80, 4771–4780

53. Gross, C., Wiesmann, V., Millen, S., Kalmer, M., Wittenberg, T., Gettemans, J., and Thoma-Kress, K. (2016) The Tax-Inducible Actin-Bundling Protein Fascin Is Crucial for Release and Cell-to-Cell Transmission of Human T-Cell Leukemia Virus Type 1 (HTLV-1). PLoS Pathog 12, e1005916

54. Gillet, N. A., Melamed, A., and Bangham, C. R. (2017) High-Throughput Mapping and Clonal Quantification of Retroviral Integration Sites. Methods Mol Biol 1582, 127–141

55. Watanabe, T. (2017) Adult T-cell leukemia: molecular basis for clonal expansion and transformation of HTLV-1-infected T cells. Blood 129, 1071–1081

56. Fogarty, K. H., Berk, S., Grigsby, I. F., Chen, Y., Mansky, L. M., and Mueller, J. D. (2014) Interrelationship between cytoplasmic retroviral Gag concentration and Gag-membrane association. J Mol Biol 426, 1611–1624

57. Roy, A., Kucukural, A., and Zhang, Y. (2010) I-TASSER: a unified platform for automated protein structure and function prediction. Nature protocols 5, 725–738

58. Yang, J., Yan, R., Roy, A., Xu, D., Poisson, J., and Zhang, Y. (2015) The I-TASSER Suite: protein structure and function prediction. Nat Methods 12, 7–8

59. Christensen, A. M., Massiah, M. A., Turner, B. G., Sundquist, W. I., and Summers, M. F. (1996) Three-dimensional structure of the HTLV-II matrix protein and comparative analysis of matrix proteins from the different classes of pathogenic human retroviruses. J. Mol. Biol. 264, 1117–1131

60. Janmey, P. A., Iida, K., Yin, H. L., and Stossel, T. P. (1987) Polyphosphoinositide micelles and polyphosphoinositide-containing vesicles dissociate endogenous gelsolin-actin complexes and promote actin assembly from the fast-growing end of actin filaments blocked by gelsolin. J. Biol. Chem. 262, 12228–12236

61. Saad, J. S., Loeliger, E., Luncsford, P., Liriano, M., Tai, J., Kim, A., Miller, J., Joshi, A., Freed, E. O., and Summers, M. F. (2007) Point mutations in the HIV-1 matrix protein turn off the myristyl switch. J Mol Biol 366, 574–585

62. Ehrlich, L. S., Fong, S., Scarlata, S., Zybarth, G., and Carter, C. (1996) Partitioning of HIV-1 Gag and Gag-related proteins to membranes. Biochemistry 35, 3933–3943

63. Scarlata, S., Ehrlich, L. S., and Carter, C. A. (1998) Membrane-Induced Alterations in HIV-1 Gag and Matrix Protein-Protein Interactions. J. Mol. Biol. 277, 161–169

64. Zhou, W., and Resh, M. D. (1996) Differential membrane binding of the human immunodeficiency virus type 1 matrix protein. J. Virol 70, 8540–8548

65. Wen, Y., Dick, R. A., Feigenson, G. W., and Vogt, V. M. (2016) Effects of Membrane Charge and Order on Membrane Binding of the Retroviral Structural Protein Gag. J Virol 90, 9518–9532

66. Doktorova, M., Heberle, F. A., Kingston, R. L., Khelashvili, G., Cuendet, M. A., Wen, Y., Katsaras, J., Feigenson, G. W., Vogt, V. M., and Dick, R. A. (2017) Cholesterol Promotes Protein Binding by Affecting Membrane Electrostatics and Solvation Properties. Biophys J 113, 2004–2015

67. Wen, Y., Feigenson, G. W., Vogt, V. M., and Dick, R. A. (2020) Mechanisms of PI(4,5)P2 Enrichment in HIV-1 Viral Membranes. J Mol Biol 432, 5343–5364

68. Bodner, C. R., Dobson, C. M., and Bax, A. (2009) Multiple tight phospholipid-binding modes of alpha-synuclein revealed by solution NMR spectroscopy. J Mol Biol 390, 775–790

69. Ceccon, A., D’Onofrio, M., Zanzoni, S., Longo, D. L., Aime, S., Molinari, H., and Assfalg, M. (2013) NMR investigation of the equilibrium partitioning of a water-soluble bile salt protein carrier to phospholipid vesicles. Proteins 81, 1776–1791

70. Hayakawa, T., Miyazaki, T., Misumi, Y., Kobayashi, M., and Fujisawa, Y. (1992) Myristoylation-dependent membrane targeting and release of the HTLV-I gag precursor, Pr53gag, in yeast. Genes & Development 119, 273–277

71. Rayne, F., Bouamr, F., Lalanne, J., and Mamoun, R. Z. (2001) The NH2-terminal domain of the human T-cell leukemia virus type 1 capsid protein is involved in particle formation. J Virol 75, 5277–5287

72. Li, L., Vorobyov, I., and Allen, T. W. (2013) The different interactions of lysine and arginine side chains with lipid membranes. J Phys Chem B 117, 11906–11920

73. Le Blanc, I., Rosenberg, A. R., and Dokhelar, M. C. (1999) Multiple functions for the basic amino acids of the human T-cell leukemia virus type 1 matrix protein in viral transmission. J Virol 73, 1860–1867

74. Behnia, R., and Munro, S. (2005) Organelle identity and the signposts for membrane traffic. Nature 438, 597–604

75. Grigsby, I. F., Zhang, W., Johnson, J. L., Fogarty, K. H., Chen, Y., Rawson, J. M., Crosby, A. J., Mueller, J. D., and Mansky, L. M. (2010) Biophysical analysis of HTLV-1 particles reveals novel insights into particle morphology and Gag stochiometry. Retrovirology 7, 75

76. Delaglio, F., Grzesiek, S., Vuister, G. W., Zhu, G., Pfeifer, J., and Bax, A. (1995) NMRPipe: A multidimensional spectral processing system based on UNIX pipes. J. Biomol. NMR 6, 277–293

77. Johnson, B. A., and Blevins, R. A. (1994) NMRview: a Computer Program for the Visualization and Analysis of NMR Data. J. Biomol. NMR 4, 603–614

78. Vranken, W. F., Boucher, W., Stevens, T. J., Fogh, R. H., Pajon, A., Llinas, M., Ulrich, E. L., Markley, J. L., Ionides, J., and Laue, E. D. (2005) The CCPN data model for NMR spectroscopy: development of a software pipeline. Proteins 59, 687–696

79. Guerry, P., and Herrmann, T. (2012) Comprehensive automation for NMR structure determination of proteins. Methods Mol Biol 831, 429–451

80. Güntert, P. (2004) Automated NMR structure calculation with CYANA. Methods Mol Biol 278, 353–378

81. Güntert, P., Mumenthaler, C., and Wüthrich, K. (1997) Torsion angle dynamics for protein structure calculations with a new program, DYANA. J. Mol. Biol. 273, 283–298

82. Keller, R. L. J. (2004) The Computer Aided Resonance Assignment Tutorial, CANTINA Verlag, Goldau, Switzerland

83. Dolinsky, T. J., Nielsen, J. E., McCammon, J. A., and Baker, N. A. (2004) PDB2PQR: an automated pipeline for the setup of Poisson-Boltzmann electrostatics calculations. Nucleic Acids Res. 32, W665–667

84. Baker, N. A., Sept, D., Joseph, S., Holst, M. J., and McCammon, J. A. (2001) Electrostatics of nanosystems: application to microtubules and the ribosome. Proc Natl Acad Sci U S A 98, 10037–10041

85. Delamarre, L., Rosenberg, A. R., Pique, C., Pham, D., and Dokhelar, M. C. (1997) A novel human T-leukemia virus type 1 cell-to-cell transmission assay permits definition of SU glycoprotein amino acids important for infectivity. J Virol 71, 259–266

